# Bacterial and viral gut microbiome alterations characterize microbiome-immune-pathophysiology axes in sickle cell disease

**DOI:** 10.64898/2026.04.13.718288

**Authors:** Zachary N. Flamholz, Jennifer De Los Santos, Karen Ireland, Janine Keenan, Jacob S. Kazmi, Aakash Mahant Mahant, Anayeli Correa, Paul S. Frenette, Betsy C. Herold, Deepa Manwani, Libusha Kelly

## Abstract

Sickle cell disease (SCD) is a chronic, inherited condition rising across the globe. Prior studies revealed a direct link between the gut microbiome and disease micropathology via aged-like (ANs) neutrophils in mouse models. In SCD patients community-level shifts in the gut microbiome included decreases in diversity and the Firmicutes/Bacteroidetes (F:B) ratio, coupled to a loss of short chain fatty acid producing microbes and a shift to non-canonical butyrate production and aerobic fatty acid oxidation pathways. ANs and the proviral microbiome associate with multiple blood cytokines, while bacterial gut microbiome features largely do not. Prophages depleted of genes related to lysis, transcriptional regulation, and host takeover were enriched in SCD patient guts, pointing to domestication of these elements, and 25% of prophages were shared at high identity between study patients. In sum, we identify prophage associated immune signatures and taxonomic and functional alterations to the gut microbiome that associate with SCD pathophysiology in a heterogeneous chronic disease both affected by and affecting microbiome composition and function.

## Introduction

Sickle cell disease (**SCD**) is a chronic, inherited hematologic disorder causing significant morbidity and mortality across the globe. SCD is characterized by pathologic red blood cell hemolysis leading to complications stemming from vaso-occlusion and vascular stress, ultimately resulting in end organ damage. In a 2018 study, it was estimated 1,950 children are born with SCD in the U.S. annually, and the life expectancy of a child with SCD is 54 years compared to 76 years for the age- and race-matched U.S. population^1^. While there has been a steady increase in life expectancy for SCD patients resulting from diagnostic screening, early intervention, and better evidence-based treatment guidelines, SCD hospitalization rates have increased^2^. Pain crises, the clinical manifestation of acute vaso-occlusion, were the number one reason for SCD-related hospital admissions from 2004-2012^2^. While gene therapy will play an important role in SCD treatment, there are significant barriers to widespread utilization^3^ and with SCD births on the rise globally^4^ alternative avenues of treatment must be pursued. In addition, novel biomarkers are valuable to better stratify SCD patients for targeted early intervention and more precise treatment decisions.

The gut microbiome is a physiological system with complex interactions with SCD. It was shown in a murine model of SCD that microbial antigens transit from the gut to the bloodstream where they activate neutrophils to a disease-promoting phenotype, termed “aged”, or “aged-like”, neutrophils (hereafter, “**ANs**”), and that depletion of gut microbes using broad-spectrum antibiotics decreased the AN population and improved inflammation-induced organ damage^5^. Human SCD patients were also found to have higher levels of the microbial antigen lipopolysaccharide in their blood and an increased AN percentage when compared to a control group of iron deficiency anemia patients with similar hemoglobin levels^6^. These findings led to the proposal of gut-targeting antibiotics as a treatment for SCD, and small studies in human subjects have shown antibiotics reduce the fraction of ANs and lipopolysaccharide in SCD patients^7–9^. The clinical momentum, though, has outpaced investigation of the SCD gut microbiome itself.

Gut microbiome studies that seek to identify microbial biomarkers of disease or disease severity look for abundance differences in individual taxa between case and control cohorts, as well as differences in summary measures related to community diversity. To date, there have been four studies that probed bacterial composition of the SCD gut microbiome compared to controls to determine if there are characteristic features of the SCD gut microbiome. Lim et al. did not observe differences in overall diversity for 35 SCD patients compared to a sickle trait control cohort, though one genus of Bacteroidetes had lower relative abundance in the SCD cohort^10^. Similarly, in a cohort of 32 pediatric SCD patients, Mohandas et al. did not observe a difference in alpha diversity or beta diversity compared to an immunocompromised cohort and a non-immunocompromised control cohort^11^. In contrast, Brim et al. observed patterns of abundance differences at all taxonomic levels below phylum when 14 SCD patients were compared to healthy controls^12^. Brim et al. also observed that SCD patients have a lower Firmicutes:Bacteroidetes (**F:B**) ratio, a gut microbiome community-level metric associated in some studies to human health^13^. Finally, in a cohort of Angolan SCD pediatric patients, Delgadinho et al. did not observe a difference in alpha diversity compared to healthy siblings but did observe abundance differences at all taxonomic ranks^14^, though fewer in number and mostly different than observed by Brim et al.

These prior studies suffer from two limitations in characterizing the gut microbiome of SCD patients. First, all studies suffered from small cohort sizes, limiting their ability to detect statistical differences in microbial taxa and population metrics. Second, the studies utilized 16S sequencing for community profiling, which cannot resolve bacterial species/strain level variation, cannot identify other microbes such as viruses, and underperforms relative to metagenomic sequencing in identifying low abundance taxa^15^. Additionally, these studies found different changes in the SCD microbiome, leading to differing conclusions about its relationship with the disease. A robust evaluation of SCD patients’ gut microbiomes, together with clinical data, is necessary to determine whether there exists a relationship between the microbial community and disease pathology.

Microbiome markers have been shown to stratify treatment response in other diseases^16,17^, providing direction for the development of clinically-meaningful gut microbiome biomarkers. Additionally, gut bacteriophage populations have been recognized for their association with chronic disease^18^ and have been shown to shift lifecycle distribution at the population level in inflammatory bowel disease^19^. Interrogation of the SCD microbiome, in conjunction with studying the effect of AN levels in SCD patients, could identify novel biomarkers of disease severity in SCD. Thus, we conducted a study of blood and stool samples from patients with SCD compared with age and race matched and otherwise baseline characteristic balanced, controls. Using whole community metagenomics, we investigated the gut microbiome on multiple axes, revealing complex yet informative patterns of microbiome change. By comparing gut microbiome signatures with blood AN and cytokine levels, we begin to characterize the pathological interplay between the gut microbiome and immune system activation in SCD.

## Results

### Sickle cell disease patient cohort

Patients and controls in the study were followed at a large hospital system in the United States. Cohorts were matched for age and race and balanced for other demographic characteristics (Supplemental Table 1). For patients, clinical history (Supplemental Table 2) and treatment history (Supplemental Table 3) were aggregated from the clinical record. Patients on prophylactic penicillin within six months of sample collection were excluded from the study due to the impact of antibiotics on gut microbiomes^21^. Additionally, laboratory measurements were collected from the clinical record by their most recent reading (Supplemental Table 4) or assayed as part of the study (Supplemental Table 5).

### Gut microbiome and virome changes in sickle cell disease

To study the SCD gut microbiome, we sequenced fecal samples from patients and controls. Sample sequencing reads were profiled for microbial taxa using a library of taxa-specific marker genes defined using shotgun metagenomic sequencing^20^. To determine whether there is a shift in the SCD gut microbiome at the community level, we used Shannon diversity to compare species diversity between patients and controls and found that SCD patients have less diverse gut microbiomes (Figure 1a). To identify microbes positively or negatively associated with SCD, we modeled taxa abundance as a function of age, sex, ethnicity, race, and SCD status using a multivariable generalized linear model (Supplemental Table 1). Overall, 25 taxa were significantly associated with SCD and 6 taxa are associated with age (Figure 1b). We noted that SCD samples had higher abundances of the phylum Bacteroidetes (**B**) and lower abundances of the phylum Firmicutes (**F**), the two taxonomic groups that make up the composite metric F:B ratio, and observe a reduction in the ratio in our study (Figure 1c), as had been observed previously^12^. Looking at human gut microbiome indicator species identified in a meta-analysis of human microbiome studies^22^, we observed a decrease in health indicators (p=5.6e-9; Mann–Whitney–Wilcoxon test) and increase in disease indicators (p=4.4e-3; Mann–Whitney–Wilcoxon test) in SCD patients compared to controls (Figure 1d), pointing to changes in the SCD gut microbiome that are consistent with changes observed in other human diseases.

**Figure 1:**
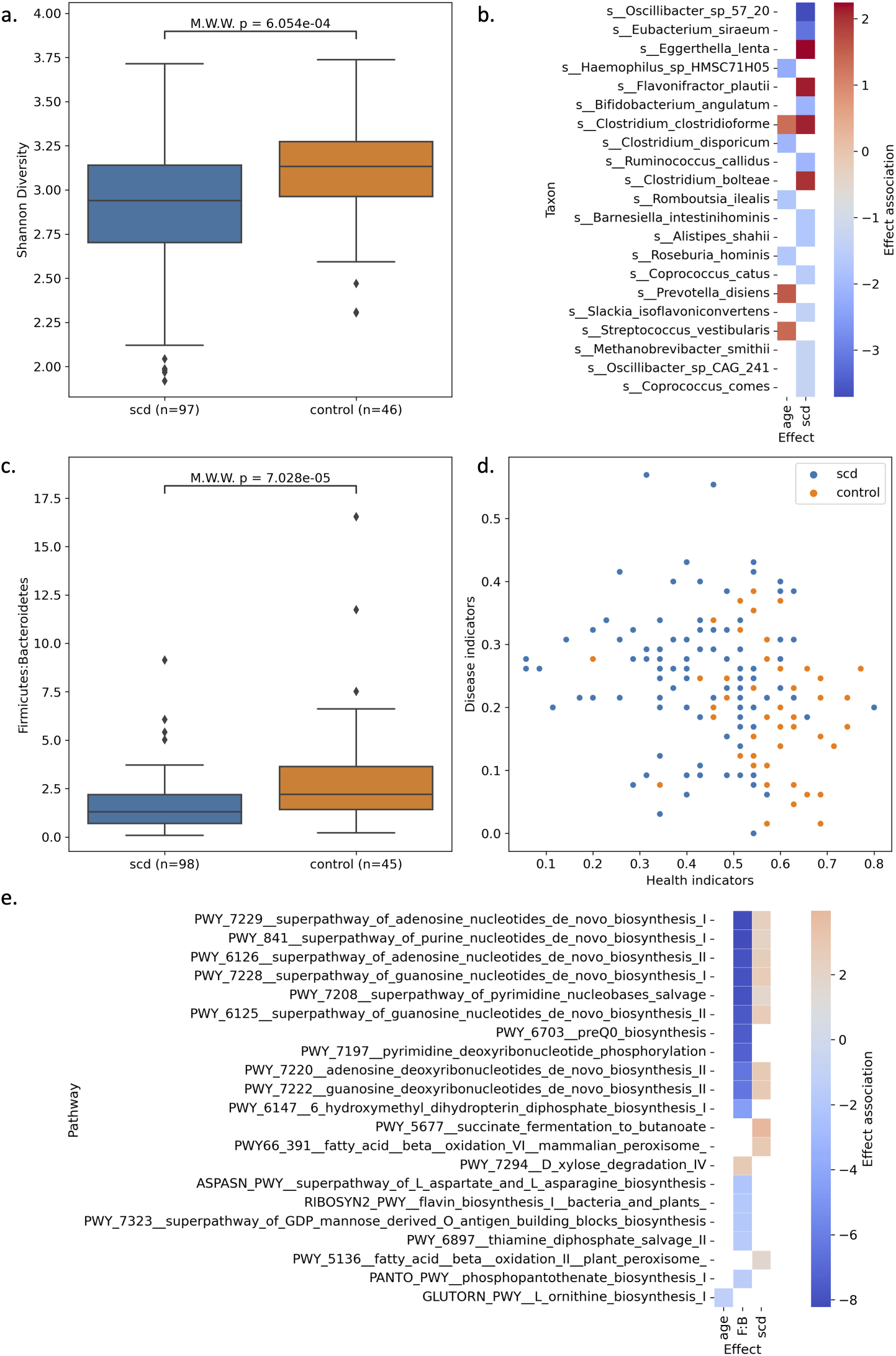
Gut microbiome community changes in SCD patients (n=98) compared to healthy controls (n=46). Sample whole community sequencing was profiled using the MetaPhlAn taxa marker database ^20^. (a) Distribution of sample alpha diversity measured as Shannon diversity for patients and controls. (b) A generalized linear model was used to determine the effect size and association of bacteria taxa abundance with sample metadata including SCD status, age, race, ethnicity, and gender, all modeled as fixed effects. Taxa with q-value *<* 0.05 for any effect are shown. (c) Distribution of Firmicutes to Bacteroidetes (F:B) ratio in the gut microbiome of patients and controls. (d) Scatter plot of health and disease indicator scores per sample. (e) Sample whole community sequencing was profiled using the HUMAnN pathway marker database ^20^. A generalized linear model was used to determine the effect size and association of pathway abundance with sample metadata including SCD status, age, race, ethnicity, and gender, all modeled as fixed effects. Additionally, the F:B was also included as fixed effect. Pathways with q-value *<* 0.05 for any effect are shown. In a and c, samples with values +/- 3 s.d. from mean of all samples were removed. M.W.W.- Mann-Whitney-Wilcoxon test. F:B, it was inversely correlated with health indicators and positively correlated with disease indicators (Supplemental Figure 1a), providing evidence that provirus enrichment mirrors changes in the bacterial species present in the sample.

We compared species composition with a beta-diversity analysis and found that beta diversity differed significantly between controls and SCD patients by Aitchison distance after CLR transformation (R^2^ = 0.030, p = 0.001). We also performed a functional characterization of samples using gene-based pathway annotation^20^. We again modeled abundance with the same fixed effects but included an additional effect, the F:B ratio, to both capture community-level changes and identify SCD-specific changes that are independent of the high-level shift (Figure 1e). High F:B ratio had exclusively negative associations that were significant, with the most significant associations with pathways related to nucleotide biosynthesis. SCD status was positively associated with three pathways related to butyrate fermentation, fatty acid oxidation, and vitamin B6 biosynthesis, and negatively associated with a pathway related to anaerobic metabolism. The small number of pathways robustly associated with SCD in our heterogeneous patient population point to specific functional changes that shed light on conserved contributions of the gut microbiome to SCD pathology.

We also interrogated the viral population in metagenomic samples. Assembled metagenomes were profiled for viral sequences using a hybrid approach that combines viral marker genes and sequence features to distinguish chromosomal, plasmid, and viral sequences^23^. Overall, there was a decrease in the number of viral sequences in SCD compared to controls (p=2.6e-5; Mann–Whitney–Wilcoxon test) (Figure 2a), likely reflecting the loss in diversity observed in SCD samples given its correlation with sample Shannon diversity and health indicators (Supplemental Figure 1a). Proviruses are lysogenic virus sequences integrated into host genomes and can be labeled in assembled viral genomes by identifying sequence features such as viral integrase proteins and integration sites^24^. Proviruses constitute a small fraction of the viral sequences present in samples, yet there was a significant enrichment of provirus sequences in SCD compared to controls (p=4.7e-4; Mann–Whitney–Wilcoxon test) (Figure 2b), which was robust to provirus or lysogenic viral sequence calling method (Supplemental Figure 2). While the provirus fraction was not correlated with bacterial community-level metrics Shannon diversity a From assembled provirus sequences it is not possible to determine whether a provirus is active, meaning replicating independently of its host via a lytic lifecycle, or inactive. Using a method developed previously^25^, we analyzed the sequencing read distribution of provirus sequences and their flanking host sequences to determine if there was increased read coverage of proviruses, which would indicate independent replication. Across SCD samples, *>*90% of proviruses were inactive (Supplemental Figure 1b), and the active provirus fraction distribution in SCD was not different from controls (p=3.208e-1; Mann–Whitney–Wilcoxon test).

**Figure 2:**
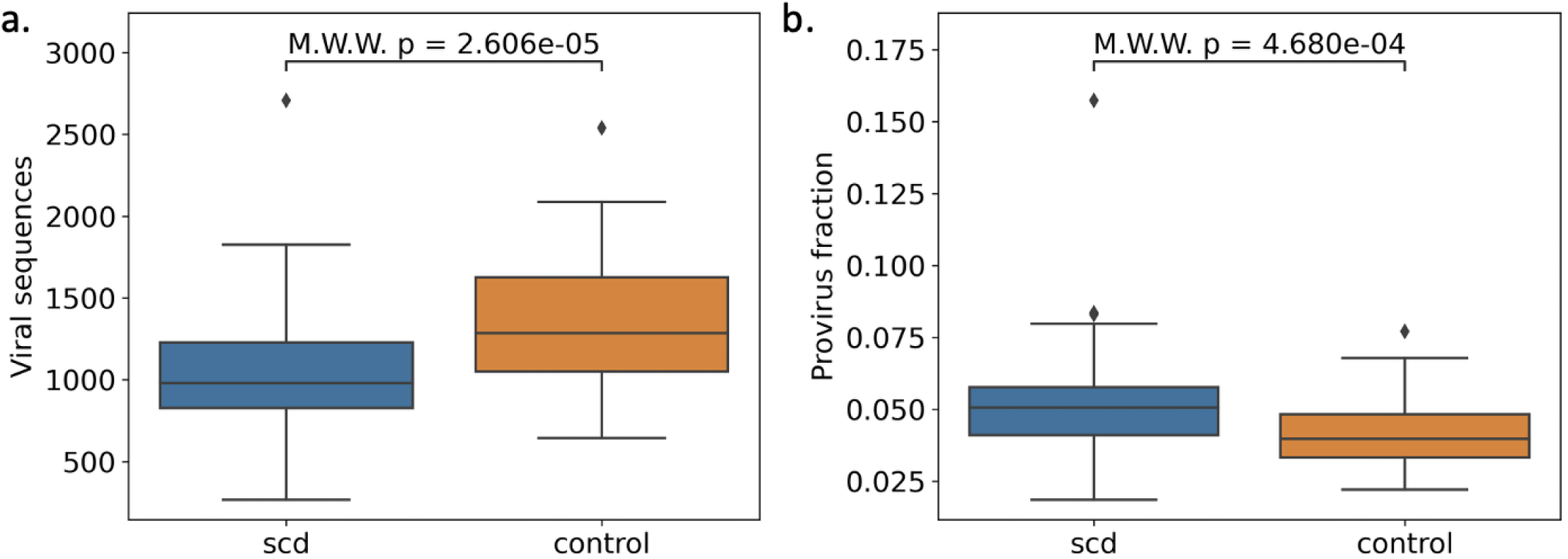
Gut microbiome viral population is altered in SCD. Sample whole community sequencing was assembled and viral sequences were identified using marker genes and sequence features. The number of viral sequences (a) and the fraction of provirus sequences (b) were compared between patients and controls. M.W.W= Mann-Whitney-Wilcoxon test.

It has been observed that SCD gut microbiomes differ significantly between both related and unrelated non-SCD patient controls^26^. Here, we asked whether control genotype, specifically sickle trait individuals (HbAS) influenced observed microbiome and virome associations. HbAS and HbAA controls did not differ significantly for F:B ratio, Shannon diversity, provirus fraction, or viral sequence count (Supplemental Table 6). In contrast to observed differences between SCD and all controls, HbAA and HbAS controls did not differ significantly in beta diversity (R^2^ = 0.024, p = 0.282), supporting the conclusion that the observed SCD/control separation was not driven by control genotype composition. We further asked whether SCD-associated microbiome and virome features were explained by clinical or demographic heterogeneity within the SCD cohort and restricted analyses to SCD participants excluding controls. No microbiome or virome feature showed an FDR-significant association with hydroxyurea use, transfusions in the past year, or acute care utilization in the past year (Supplemental Table 7).

### Gut microbiome and virome markers and known SCD biomarkers

We investigated whether gut microbiome changes in SCD are related to the AN population by asking whether microbiome markers correlate with the fraction of ANs of the total neutrophil population in a patient’s blood. For this analysis, we utilized a subset of the SCD cohort whose blood was assayed (n=57). Controls were not assayed for ANs and were therefore excluded from the following analysis. We evaluated four groups of markers: (1) species positively associated with SCD- *E. lenta*, *F. plautii*, and *C. bolteae*; (2) species negatively associated with SCD- *O. sp. 57 20*, *E. siraeum*, *A. shahii*, *B. intestinihominis*, *R. callidus*, *B. angulatum*, *C. catus*, *S. isoflavoniconvertens*, *M. smithii*, *O. sp CAG 241*, and *C. comes*; (3) functions significantly enriched between patients and controls-succinate fermentation to butanoate, fatty acid beta oxidation VI mammalian peroxisome, and fatty acid beta oxidation II; and (4) system-level summary metrics shown to be differential between patients and controls-Shannon diversity, F:B ratio, health indicators, disease indicators, viral counts, and provirus fraction. A single marker, the bacterial species *A. shahii*, had a nominally significant correlation with the AN percent (*ρ* = 0.26, p=0.048).

Given the substantial changes in the gut microbiome of SCD patients, we next investigated whether gut microbiome features are associated with SCD clinical and molecular measures of disease severity. We utilized a molecular blood panel to profile SCD patients for levels of cytokines and chemokines (Supplemental Table 5). Additionally, we included a number of hemolysis-related and disease burden clinical measurements (Supplemental Table 4).

We first evaluated the AN fraction given its known role in SCD pathology. Neutrophil activation assay measures had significant correlations with white blood cell measures collected from the health record (Supplemental Figure 3), providing confidence in comparing AN data with clinical data. ANs were significantly correlated with four inflammatory cytokines: interferon gamma, interleukin-1*β*, interleukin-10, and interleukin-17A, and one chemokine: monocyte chemoattractant protein-1 (Figure 3, top).

**Figure 3:**
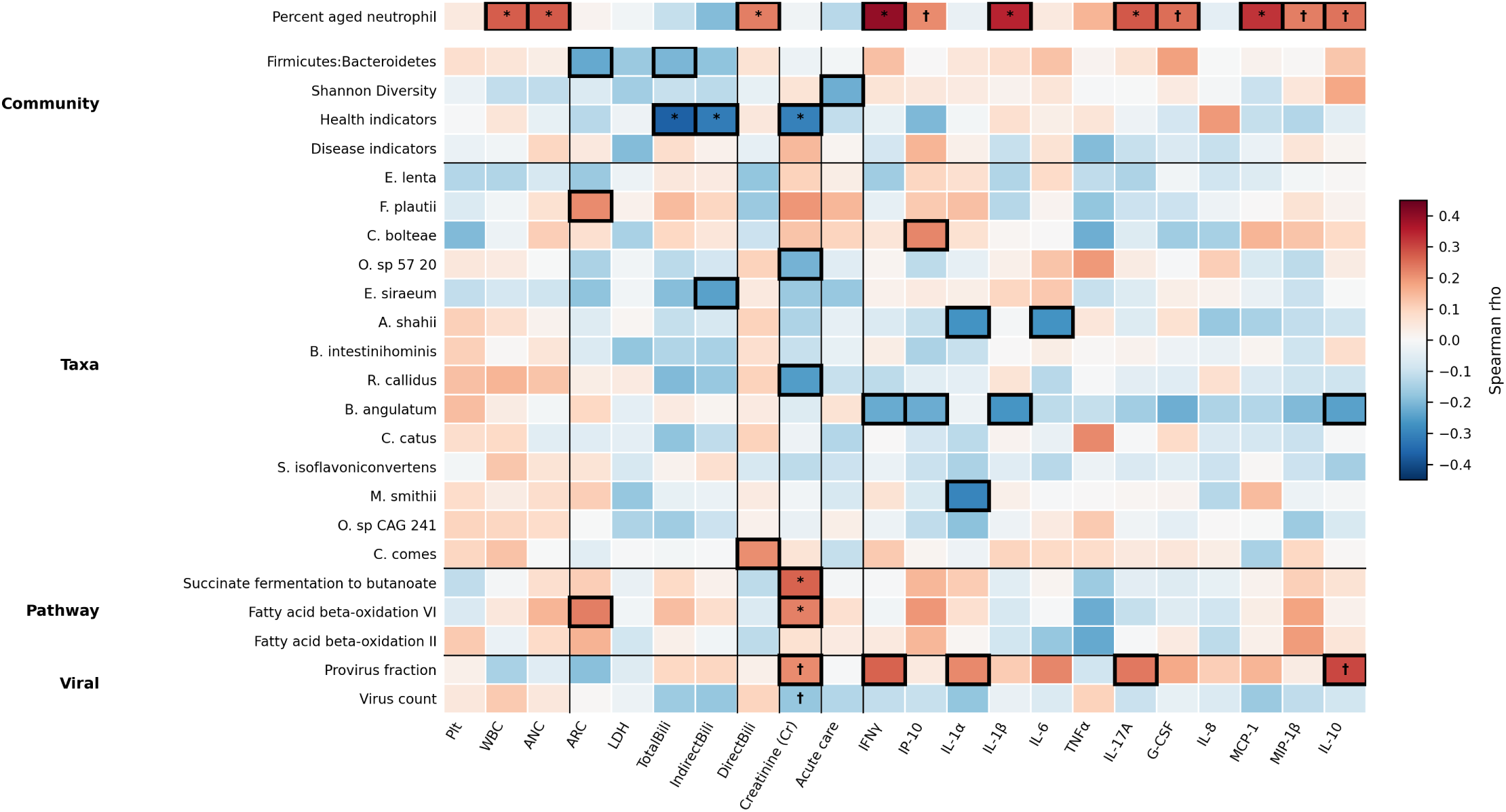
Aged neutrophil percentage (n=57) and gut microbiome and virome feature (n=98) correlations with clinical measures and blood molecular immune markers. Correlations were measured with Spearman *ρ* and significance was measured using permutation testing. Rows are grouped as community metrics, taxa, functional pathways, and viral features. Columns are grouped as hematologic/inflammatory markers, hemolysis markers, liver marker, kidney marker, acute care burden, and cytokines/chemokines. Boxes denote nominal p *<* 0.05; * denotes FDR q *<* 0.05; † denotes FDR q *<* 0.10. FDR correction was applied within each feature-group by marker-block family. For the number of samples assayed for each clinical and molecular measure, see Supplemental Tables 4-5.

In SCD-only correlation analyses, microbiome and virome features were tested against clinical markers and cytokines using Spearman correlations, with FDR correction applied within prespecified feature-group by marker-group blocks (Figure 3, bottom). Lower health-associated microbiome indicator scores were significantly associated with higher total bilirubin, indirect bilirubin, and creatinine. Functional pathway features, including succinate fermentation to butanoate and fatty acid beta-oxidation VI, were also positively associated with creatinine. In contrast, the viral features, provirus fraction and virus count, did not show broad FDR-significant associations with clinical markers after block-level correction. These analyses suggest that bacterial community and functional features are most strongly related to hemolysis- and renal-associated clinical variation, while provirus fraction shows a more consistent immune association and is significantly associated with interleukin-10.

### Integrated viruses

The provirus fraction in the gut microbiome had correlation trends with multiple inflammatory cytokines in SCD patients. A total of 4,962 proviruses were identified across the SCD patient gut microbiomes. Provirus lengths had a bimodal distribution with a median size of 23,775 base pairs (Supplemental Figure 4). All proviruses with predicted taxonomy (99.5%) were bacteriophages of the class Caudoviricetes (99.3%), meaning the proviral sequences are prophages.

Because the vast majority of phages were categorized as inactive in host bacterial genomes, we wondered whether these sequences would be similar between patients. We clustered all prophage sequences at 99% identity over at least 70% bi-directional coverage and found 395 clusters of sequences with median size of 2 (Figure 4a). In total, 1,249 (25%) sequences had a nearly identical sequence in another sample in the dataset. These clustered prophages were homologous to sequences in common gut commensals, however Bacteroides stood out with 113 clusters matching species in the genus (Supplemental Figure 5). The decreased F:B ratio in our patient population may also represent an increase in Bacteroides strains carrying these conserved prophages, although at the community level, the prophage fraction did not inversely correlate with the F:B ratio as would be expected if it were such strains driving the community shift. To determine whether clustered prophages differed from singletons, we compared the high-level functional content of sequences in each group (Figure 4b). Interestingly, clustered prophage sequences had lower fractions of genes with functions related to lysis, transcriptional regulation, and host takeover and a higher fraction of genes with unknown function compared to non-clustered prophages. A reduction in proteins that function in host lysis was also observed in active prophages compared to dormant prophages (Supplemental Figure 6).

**Figure 4:**
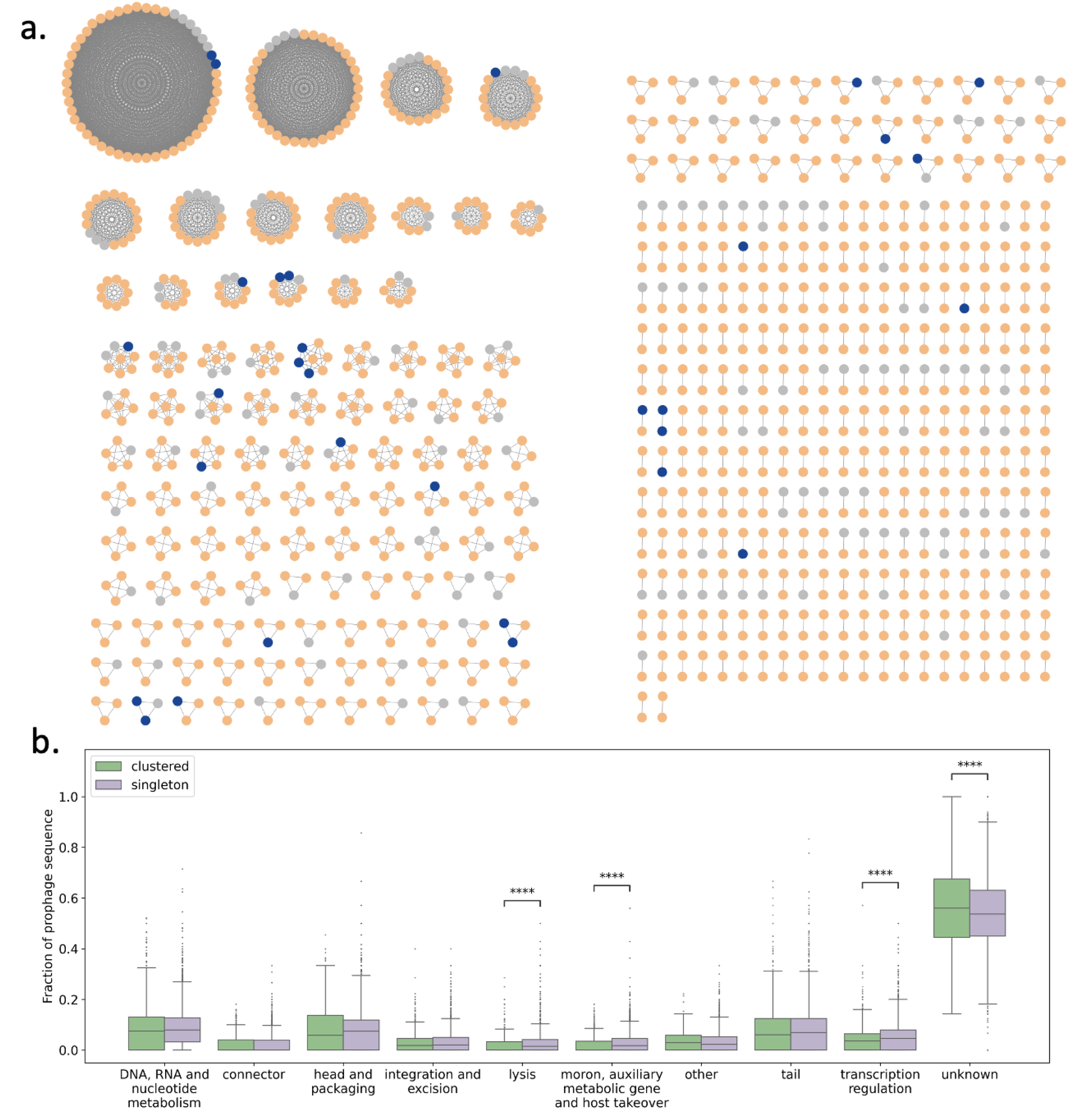
Highly conserved prophage sequences are shared by gut microbiomes of SCD patients. (a) Patient microbiomes sharing a highly conserved prophage sequence visualized as fully connected networks. Conserved prophages had 99% sequence identity over at least 70% of bidirectional sequence coverage. Node color indicates predicted prophage activity: active (blue), dormant (orange), and undetermined (gray). (b) Prophage sequence distribution of high-level viral protein functions compared between clustered (n=1,249) and singleton (n=3,713) prophages. Significance tested with a Mann-Whitney-Wilcoxon test: **** = p *<* 0.0001.

## Discussion

The primary goal of our study was to ask whether there exist consistent, significant interactions between the gut microbiome and disease pathology in SCD. Such interactions, when observed across heterogeneous patients with varying clinical histories and burdens of disease, could lead to novel treatment modalities and diagnostics to improve patient care and monitoring in this disease of increasing importance around the world. With orthogonal computational and experimental approaches, we can begin to build a model of the interactions between gut microbiome features, immune system activity, and pathology in SCD.

Our study reveals a structured dialogue between the gut ecosystem and systemic inflammation in SCD (Figure 5). Shotgun metagenomics of 98 patients showed a disease-wide contraction of bacterial diversity, a drop in the F:B ratio, and characteristic changes in pan-disease indicator taxa. These shifts parallel patterns seen across disparate chronic illnesses^22,27^, suggesting that SCD is associated with a broader disease-associated microbiome signature at the community level.

**Figure 5:**
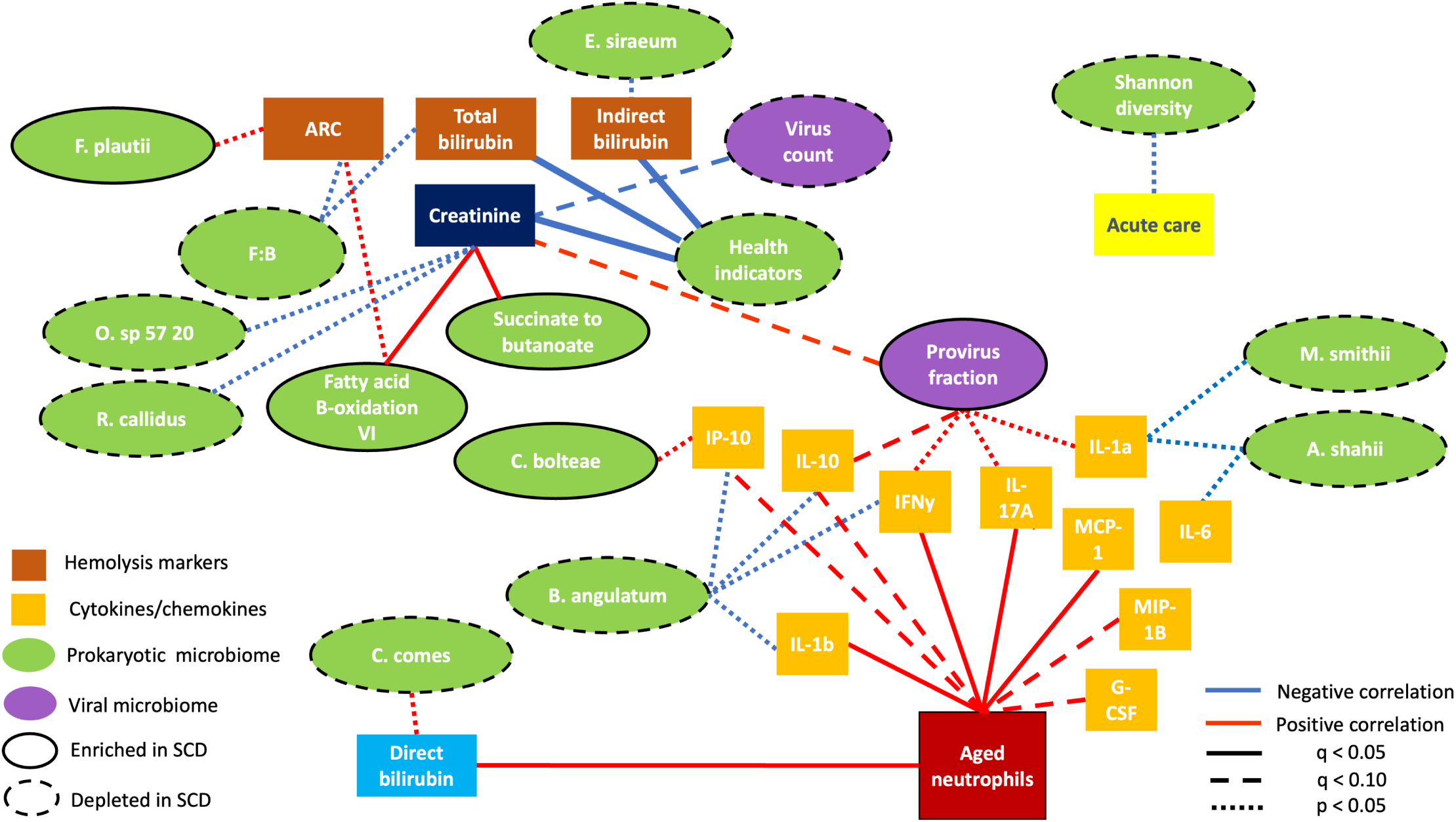
A model of interactions between the gut microbiome and markers of inflammation in the blood of SCD patients. Using multiple data modalities, we begin to understand the complex interaction of gut microbiome changes and immune system activation in the pathology of SCD. Clinical and molecular features are shown as boxes, microbiome and virome features as ovals. Edges represent associations and do not represent causal or mechanistic interactions. Edge style indicates significance: dotted line (nominal p *<* 0.05), dashed line (FDR q *<* 0.10), solid line (FDR q *<* 0.05)

Neutrophil biology and the microbiome appear to occupy distinct, complementary niches. The AN subset correlated with an overlapping cytokine panel but less with bacterial community metrics. Conversely, bacterial taxa and health indicator taxa associated with indirect bilirubin, total bilirubin, and reticulocyte count, clinical proxies for hemolysis, yet show little relationship to the immune milieu. This dissociation implies that neutrophil activation and microbial changes represent parallel arms of SCD pathophysiology, informative for different disease facets. We observe multiple associations, both nominal and significant, between microbiome taxonomic and functional features, and hemolysis markers. Health indicators are significantly negatively associated with total and indirect bilirubin, F:B ratio is nominally negatively associated with ARC and total bilirubin, the SCD-depleted taxa *E.siraeum* is nominally negatively associated with indirect bilirubin, and the SCD-enriched taxa *F.plautii* is nominally positively associated with ARC. We observe negative associations between creatinine and microbiome health indicators, SCD-depleted bacteria, and virus count. We observe positive associations between creatinine and SCD microbiome functions related to fatty acid B-oxidation and succinate to butanoate fermentation, and provirus fraction. Associations between serum creatinine and gut microbiome features have been observed in prospective study of chronic kidney disease^28^. We note that SCD patients frequently suffer from nephropathies^29^; the associations we observe here require confirmation in future studies with additional measures of renal function.

Dormant prophages emerge as a previously unrecognized feature of the SCD gut microbiome associated with immune variation. We detect a 1.7-fold increase in the prophage fraction, yet *>*90% of these elements are predicted to be replicatively silent. Strikingly, the proportion of prophages, but not total phage population or lytic phage fraction, tracks with IL-1 α, IL-10, IL-17A and IFN-γ levels. Immune signaling molecules are known to be elevated at baseline in SCD patients^30–32^, and 3/4 cytokines correlated with prophage level are observed to be lower in patients in vaso-occlusive crisis compared to SCD patients not in vaso-occlusive crisis, including IL-10, IL-17A, IFN*γ* ^30^. Interestingly, the gut microbiome in SCD patients harbors prophage sequences that are highly conserved between unrelated individuals, with one such sequence present in over 50% of metagenomes. These conserved, shared prophages were an unexpected feature of SCD patient microbiomes, and stand in contrast to studies of lytic phages, that found non-integrating bacteriophage populations are highly individual-specific ^33^. We found that clustered sequences had a lower fraction of genes coding for proteins involved in host takeover, gene expression manipulation, and lysis. We speculate that lysogens bearing ‘domesticated’ prophages maintain a survival advantage in the setting of a disease-associated gut microbiome. Enrichment of lysogens may modulate host immunity by altering bacterial surface antigens or metabolite flux. Experimental induction assays and longitudinal sampling will be needed to test whether prophage excision precedes cytokine spikes or merely registers host stress, and a deeper analysis of shared prophage host range, transmission, and association with other microbiome-influencing variables is required to assess if shared prophages are specific to SCD biology or a more general phenomenon.

Functional signature investigation of the gut microbiome, enabled by metagenomics, can have therapeutic implications^34^. Among the functional module changes observed in SCD, three merit particular attention: (i) enrichment of nucleotide biosynthesis pathways, (ii) succinate to butanoate fermentation, a non-canonical route for microbial butyrate generation, and (iii) fatty-acid *β*-oxidation pathways, which typically require molecular oxygen and are not common in the anaerobic gut environment. The communal shift toward a low F:B ratio was tightly coupled to an expansion of nucleotide-biosynthesis modules, offering a narrow-spectrum antimicrobial target set that could re-balance the ecosystem more precisely than broad antibiotics trialed to date^7–9^. Looking at functional modules enriched in SCD independent of community shift, we see the succinate to butanoate butyrate production pathway; a departure from the primary gut butyrate acetyl-CoA pathway^35^. Butyrate is the primary energy source for gut epithelial cells and has been shown to be anti-inflammatory and supportive of epithelial-barrier integrity^36^. This shift in butyrate production pathway in the SCD gut may reflect the loss of members of taxonomic groups that produce butyrate via complex carbohydrate fermentation, such as *C. catus*, *C. comes* and *E. siraeum* ^35,37^. We note that previous SCD rodent model work observed decreased butyrate in disease^38^; further work is necessary to understand the interaction between different mechanisms of butyrate production in the SCD microbiome. Conversely, elevated *β*-oxidation genes suggest greater flux through short- and medium-chain fatty-acid catabolism, a shift that might deplete beneficial metabolites and warrants metabolic follow-up. Notably, the SCD enriched *β*-oxidation pathways II and IV are mitochondrial and peroxisomal, and therefore generally aerobic, pointing to the availability of molecular oxygen in the gut, a hallmark of gut barrier disruption ^39,40^.

There are a number of recognized limitations for our study. First, while we collectively grouped SCD patients into a single cohort, the disease can vary in its presentation and clinical course, which may affect the gut microbiome. Our cross-sectional design precludes causal inference and we acknowledge that as a chronic disease, SCD presentation can be highly variable. Second, flow-cytometry assays for neutrophil activation markers were performed within 8 hours of venipuncture, and while we have correlative evidence that our results reflect accurate measures, we did not formally benchmark marker stability to this time window. Next, prophage activity was in-ferred from coverage ratios, as is common in the field, but with this method asynchronous induction events could evade detection. Finally, findings derive from a single U.S. center cross-sectional study and may not generalize to global SCD populations with different diets or genotypes. We note that F:B ratio is a non-specific, but commonly used microbiome metric that is subject to confounding.

Due to patient heterogeneity, individual associations are difficult to detect, limiting the generalizability of observations made in this study. Yet, consistent patterns across related biomarkers may still be informative. Thus, we see utility in considering the directionality of microbiome and virome associations in addition to the statistical significance of individual outcomes. As an example, B. angulatum, a bacterial taxon depleted in SCD, showed the most consistent nominal associations with immune markers among the taxa, despite none surviving FDR correction. Higher-level microbiome and virome features are limited, but also useful because they may capture shared ecological or functional states. For example, virus count shows an almost opposite pattern from prophage fraction across many clinical and molecular measures, although individual virus count associations generally do not reach nominal significance. More broadly, features depleted in SCD microbiomes, including specific taxa, health indicators, and virus count, tend to associate with both clinical pathophysiology and molecular markers of immune activation despite most associations being nominal only. These patterns do not establish causality or identify specific mechanisms, but they begin to define a landscape of positive and negative associations that support unified features of SCD microbiomes and can guide future mechanistic studies.

In summary, we propose that the gut microbiome bacterial shift mirrors hemolytic burden in SCD while ANs and dormant prophages more strongly mirror immunological activation. Longitudinal and interventional studies, particularly those integrating metatranscriptomics and targeted bacteriotherapy, are necessary to unravel causality and mechanism and potentially translate these observations into clinical tools.

## Declarations

### Ethics approval and consent to participate

This study was approved by the Institutional Review Board of the Albert Einstein College of Medicine and Montefiore Medical Center (IRB 2018-9080). All study participants were recruited from the outpatient clinics at the Children’s Hospital at Montefiore and Montefiore Medical Center. Written informed consent was obtained from all participants or their legal guardians prior to enrollment.

### Consent for publication

Not applicable.

### Data availability

The dataset “Sickle cell disease patient gut microbiome study” is available in the repository NCBI and can be accessed via the following accession ID: PRJNA1320713. Patient level clinical, neutrophil, and immune data can be found in the Supplementary Material file ‘SCD study data.xlsx’.

### Authors’ contributions

Z.N.F. developed the hypotheses, analyzed the data, and prepared all figures. Z.N.F and L.K. wrote and revised the manuscript text with D.M. and B.C.H contributing. D.M. and P.S.F. designed the study and secured IRB approval. J.D.L.S., K.I., and J.K. supported study participant enrollment, sample collection, and data aggregation. J.S.K. conducted the neutrophil assay. A.M.M. and A.C. conducted the immune profiling assays.

## Acknowledgments

The authors thank the reviewers for their constructive comments that greatly improved the manuscript. The authors thank Marcel R. M. van den Brink and Jonathan U. Peled at the Molecular Microbiology Facility of Memorial Sloan Kettering Cancer Center for their generous assistance with microbiome sequencing. The authors thank the Kelly and Herold labs for helpful discussions.

## Methods

### Study design and subjects

The study is a single-center outpatient cohort study of SCD patients and age- and race-matched controls. Study participants were recruited from the Children’s Hospital at Montefiore and Montefiore Medical Center (IRB NUMBER: 2018-9080). Patients were enrolled with the following inclusion criteria: 1) diagnosis of sickle cell anemia (sickle cell disease SS genotype or sickle beta zero thalassemia, S*β*^0^-thal), or sickle cell trait in association with hereditary persistence of fetal hemoglobin (S-HPFH); 2) age ≥ 4; and 3) currently in a usual state of health. Exclusion criteria included: 1) malignancy; 2) inflammatory bowel disease or other gastrointestinal disorder; 3) immunocompromised due to additional underlying disease or immunosuppressive medication; 4) history of hematopoietic stem cell transplant; 5) history of C. difficile infection in the preceding 2 months; 6) receipt of chemotherapy in the preceding 2 months; 7) receipt of systemic antibiotic other than Pen VK in the preceding 2 months; 8) Hospitalization within the preceding 2 weeks; 9) intercurrent febrile illness or sickle cell related pain episode requiring opioids within the preceding 2 weeks. Controls were included with age ≥ 4 and excluded with the same criteria in addition to any systemic antibiotic use in the preceding 2 months.

In total, 101 SCD patients and 66 controls were enrolled in the study from 2018-2020. Of these participants, 98 SCD patients and 46 controls provided stool samples. One patient sample did not have enough material for metagenomic sequencing. Fecal samples were collected by study participants into DNA/RNA Shield Fecal Collection tube (Zymo Research) and samples were split into 6 tubes and stored at -80*^◦^*C. Blood was also drawn from study participants around the time of fecal collection.

### Clinical and molecular data

Clinical data was collected for SCD patients based on chart review. Information included: (i) demographics-age (years), sex, ethnicity, race, hemoglobin genotype (SS, S*β*^0^, SS with high F, AS, AA, where ‘S’ indicates a sickle cell allele, ‘*β*^0^’ indicates a thalassemia allele coding for no wild type hemoglobin, ‘F’ indicates a fetal hemoglobin allele, and ‘A’ indicates a wild-type allele); (ii) patient history-hydroxyurea (on/off), folic acid (on/off), glutamine (on/off), stroke (yes/no), asthma (yes/no), allergies (yes/no), bacteremia (yes/no), silent infarct (yes/no), history of meningitis, osteomyelitis, or urinary tract infection (yes/no), number of acute chest syndrome past year, number of pain admissions past year, number of 30 day readmissions past year, number of emergency room visits past year, number of transfusions in the past year, number of exchanged transfusions in the past year; and (iii) clinical measurements-weight (kg), body mass index (BMI), urine microalbumin (MA/Creatinine; mg/gm), lactate dehydrogenase (LDH; U/L), platelet (PLT; k/uL), white blood cells (WBC; k/uL), absolute neutrophil count (ANC; k/uL), hemoglobin (Hb; g/dL), sickle hemoglobin (HbS; %), fetal hemoglobin (HbF; %), wild type hemoglobin (HbA; %), absolute reticulocyte count (ARC; k/uL), total bilirubin (mg/dL), direct bilirubin (mg/dL), indirect bilirubin (mg/dL), alanine aminotransferase (ALT; U/L), Creatinine (mg/dL), iron studies: iron level (ug/dL), transferrin (ug/dL), saturation (%), total iron binding capacity (TIBC; ug/dL), ferritin (ng/dL). All yearly measures were annualized to 1 year if patient records are *<* 1 year in duration. Acute care in the past year is the sum of pain admissions and emergency department visits.

Venous blood in Acid Citrate Dextrose was collected from study participants. Inflammatory cytokine and chemokine levels were measured from serum using the MILLIPLEX Human Cytokine/Chemokine/Growth Factor Panel A kit (MilliporeSigma, HCYTA-60K). Luminex assay data were acquired on a Luminex MAGPIX and analyzed with the MILLIPLEX Analyst program (MilliporeSigma). The following cytokines and chemokines were profiled: G-CSF (pg/mL), IFNγ (pg/mL), IL-10 (pg/mL), IL-17A (pg/mL), IL-1α (pg/mL), IL-1*beta* (pg/mL), IL-6 (pg/mL), IL-8 (pg/mL), IP-10 (pg/mL), MCP-1 (pg/mL), MIP-1*β* (pg/mL), TNF-α (pg/mL). Cytokines and chemokines with *<* 50% of samples assayed at the lower limit of detection were excluded from further analysis.

### Clinical and immune marker organization

Clinical and immune markers were organized into prespecified marker groups: hematologic/inflammatory markers, including platelet count, WBC, and ANC; hemolysis markers, including ARC, LDH, total bilirubin, and indirect bilirubin; a liver-associated marker, direct bilirubin; a renal marker, creatinine; acute care utilization in the past year; and individual cytokines/chemokines, including IFNγ, IP-10, IL-1α, IL-1*beta*, IL-6, TNF-α, IL-17A, G-CSF, IL-8, MCP-1, MIP-1*β*, and IL-10. We recognize that some markers could belong in multiple categories.

### Neutrophil activation biomarker assessment

Blood was also evaluated by flow cytometry for neutrophil adhesion and activation markers. Flow cytometry was performed using LSRII equipped with FACS Diva 8.0.1 software (BD Biosciences) and analyzed with FlowJo software (Tree Star). Neutrophils were identified by forward and side scatter characteristics combined with CD16b expression, and the AN subset evaluated by CD62L^lo^ CXCR4^hi^ expression within the neutrophil population. Whole blood from the same sample used for flow cytometry assays was diluted 1:10 in PBS for complete blood count with differential counts on ADIVA 120 (Siemens Healthcare Diagnostics). Total WBC and absolute neutrophil counts along with percent AN from flow cytometry were used to calculate the absolute aged neutrophil count. Samples were excluded if analyses were not performed within 8 hours of blood draw.

### DNA extraction and sequencing

Sample preparation and sequencing was done by the Molecular Microbiology Facility of the Integrated Genomics Operation at Memorial Sloan Kettering Cancer Center. DNA was extracted from samples using a custom phenol chloroform extraction optimized for fungal and bacterial isolation. Metagenomic sequencing was performed using next generation sequencing with the Illumina HiSeq platform using 2×150bp paired end reads. For some samples, multiple sequencing runs were performed to achieve targeted sequencing depth. Base calling was performed by platform software resulting in paired FASTQ files for each sample. Samples with multiple runs were concatenated to single paired files.

### Metagenomic sequence profiling

For bacterial taxa and function analysis, the bioBakery 3 suite of metagenomic sequence tools was used ^20^. First, raw reads were fist processed for quality and removal of human contamination using kneaddata (v0.10.0) with the following parameters -t 8 -p 12 –cat-final-output^20^. Bacterial taxa were profiled with MetaPhlAn3 (v3.0) with default parameters and –no map –nproc 12 and utilized the mpa v30 CHOCOPhlAn 201901 clade-specific marker gene database^20^. MetaPhlAn output for all samples were collapsed to a single table using the merge metaphlan tables.py utility script^20^. Function pathways were profiled with HUMAnN3 (v3.7) ^20^. Paired read files were merged to a single table and HUMAnN was run with default parameters and –threads 96^20^ and database versions mpa vOct22 CHOCOPhlAnSGB 202212 and uniref90 201901b full.dmnd. For comparison between samples, pathway profiles were normalized for read sequence depth using the script humann renorm table to copies per million with parameters –units cpm –update-snames; stratified profiles were split using the humann split stratified table script; and merged to a single table using the humann join tables script^20^.

For viral profiling, raw sequences were processed with the nextflow nf-core/mag (v2.4.0) metagenomics pipeline^41,42^ for quality and contamination removal, assembly with MEGAHIT ^43^, and gene calling with prodigal^44^. The pipeline was run with -profile singularity and parameters –skip binning –skip spades. Viral sequences were profiled from MEGAHIT assemblies using geNomad (v1.7.1) with the end-to-end command and default parameters^23^ and using VIBRANT (v1.2.1) using the VIBRANT run.py script with -f nucl -t 8 -no plot parameters^45^. Both methods predict provirus sequences but only VIBRANT predicts lytic vs. lysogenic bacteriophage life cycle. To calculate the non-integrated lysogenic virus count in the VIBRANT data, the count of provirus sequences was subtracted from the count of lysogenic viruses per sample. PropagAte (v1.1.0) with default parameters was used to determine the activity of provirus sequences ^25^ from raw sequence reads, MEGAHIT assemblies, and provirus coordinates called by both geNomad and VIBRANT. All fraction metrics were calculated by dividing the count of interest by the total number of viruses called by the respective method (e.g. the provirus fraction for geNomad is the count of proviruses divided by the total count of viruses per sample).

### SCD metagenome analysis

Multiple metrics were calculated for the bacterial population in each sample using its MetaPhlAn profile. Shannon diversity was calculated using the MetaPhlAn utility script calculate_diversity.R script with parameters -d alpha -m shannon -s s_.The F:B ratio was calculated by dividing the abundance of ‘t_Firmicutes’ by the abundance of ‘t_Bacteroidetes. Health and disease associated taxa were collected from a gut microbiome meta-analysis^22^, and the number of taxa in each set was counted based on the presence of the taxa at any abundance in the sample, as performed in^22^.

To determine bacterial taxa and pathways differentially abundant in SCD patient samples compared to controls, we utilized a multivariate generalized linear model approach to model profiles as log-linear with MaAsLin 2 (R; v1.8) ^46^. For taxa, MaAslin2 was run with min prevalence = 0.1 and min abundance = 0.0001, using MaAsLin2 default total-sum scaling normalization and log transformation. For pathways, MaAslin2 was run with min prevalence = 0.1 and normalization = ‘NONE’. We included a number demographic variables as fixed effects. We modeled feature abundance as:

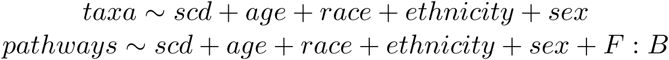

where scd is an indicator variable for disease status and abundance refers to either taxonomic or pathway relative abundance. Because metagenomic relative abundance profiles are compositional, MaAsLin2 coefficients were interpreted as associations with transformed relative abundance rather than absolute microbial abundance. Effect heat maps were generated from MaAslin2 significant result tables, and significant effects were reported for q*<*0.05.

### Prophages investigation

Prophage sequences identified using geNomad were clustered at 99% sequence identity over *>*= 70% of bidirectional sequence coverage using mmseqs2 (v14.7e284)^47^ with parameters –cluster-mode 0 –cov-mode 0 -c 0.7 –min-seq-id 0.99. Clusters were converted to fully connected networks using the python package networkx (v3.1)^48^ and visualized using cytoscape (v3.10.0)^49^. VPF-PLM^50^ was used for functional annotation of prophage sequences. Clustered prophage homology to bacterial species sequences was done using blastn (v2.16.0+) and the core nt database with parameters evalue 1e-10 and perc identity 90. The best hit bacterial species was assigned to each cluster where such a hit was identified.

### Beta diversity analyses

We conducted a beta diversity analysis using MetaPhlAn species profiles. For the primary disease/control comparison, samples were grouped as control or SCD. For the genotype control sensitivity analysis, samples were restricted to HbAA and HbAS individuals. Species detected in at least 10% of included samples were retained. To account for the compositional structure of metagenomic relative abundance data, species profiles were transformed using a centered log-ratio transformation after addition of a small pseudocount to accommodate zero values. Aitchison distances were calculated from the CLR-transformed species profiles. Statistical significance of group separation was assessed by PERMANOVA using 999 permutations. For the control versus SCD comparison, PERMANOVA was performed between the two disease-status groups. For the HbAA versus HbAS control comparison, PERMANOVA was performed among controls only. Analyses were implemented in python using pandas^51^ and NumPy^52^ for data processing, scikit-bio for distance calculations and PERMANOVA, scikit-learn^53^ for ordination, and matplotlib^54^ for visualization.

### Statistical analyses and visualizations

To test whether SCD-associated microbiome and virome features were explained by clinical or demographic heterogeneity within the SCD cohort, we restricted analyses to SCD participants, excluding controls. We fit a separate multivariable regression model for each feature. Each model included age, sex, hydroxyurea use, transfusions in the past year, and acute care utilization in the past year as predictors.

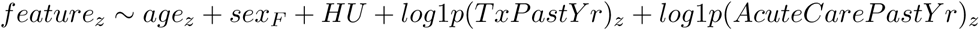

Non-negative abundance, ratio, pathway, viral, and count-like variables were log-transformed using feature-specific pseudocounts for zero-containing microbiome/virome variables and ln(1 + x) transformation for count covariates. Diversity and indicator scores were not log-transformed. Continuous variables were standardized to Z-scores before modeling. Models were fit using ordinary least squares with HC3 robust standard errors. FDR correction was applied separately for each model term across the tested microbiome and virome features.

Mann–Whitney–Wilcoxon was used to compare distribution means between SCD patients and controls for Shannon diversity, F:B ratio, healthy indicator fraction, and disease indicator fraction. Values were removed in +/- 3 standard deviations from mean of all study samples. For flow cytometry and electronic health record white blood cell measures, correlation was tested with Pearson correlation coefficient. Mann–Whitney–Wilcoxon test was used for all tested association of categorical variables test with a two-sided alternative hypothesis. All methods were implemented with scipy stats package^55^: spearmanr, pearsonr, and mannwhitneyu with default parameters and permutation test with alternative=’two-sided’, permutation type=’pairings’. Visualizations were produced using python packages seaborn^56^ and statannotations^57^. Pandas^51^ and numpy^52^ python packages were used for analysis.

To test relationships between microbiome/virome features and clinical or immune markers within the SCD cohort, we performed Spearman correlation analyses using the main SCD cohort only, excluding controls. Microbiome and virome features were organized into four prespecified feature groups: community metrics, taxa, functional pathways, and viral features. Clinical and immune markers were grouped into marker sets for visualization and multiple-testing correction, including hematologic/inflammatory clinical markers, hemolysis markers, creatinine, acute care burden, and individual cytokines/chemokines. Spearman correlation coefficients were calculated for each feature–marker pair. Benjamini-Hochberg FDR correction was applied within each prespecified feature group by marker group block. Nominal associations were defined as p *<* 0.05, FDR-significant associations as q *<* 0.05, and trends as q *<* 0.10. In the heatmap, boxes indicate nominal p *<* 0.05, asterisks indicate q *<* 0.05, and daggers indicate q *<* 0.10. The resulting data describes pairwise relationships and does not test that each association is independent of age, sex, treatment, disease severity or other covariates.

## Supplemental

**Supplemental Figure 1:**
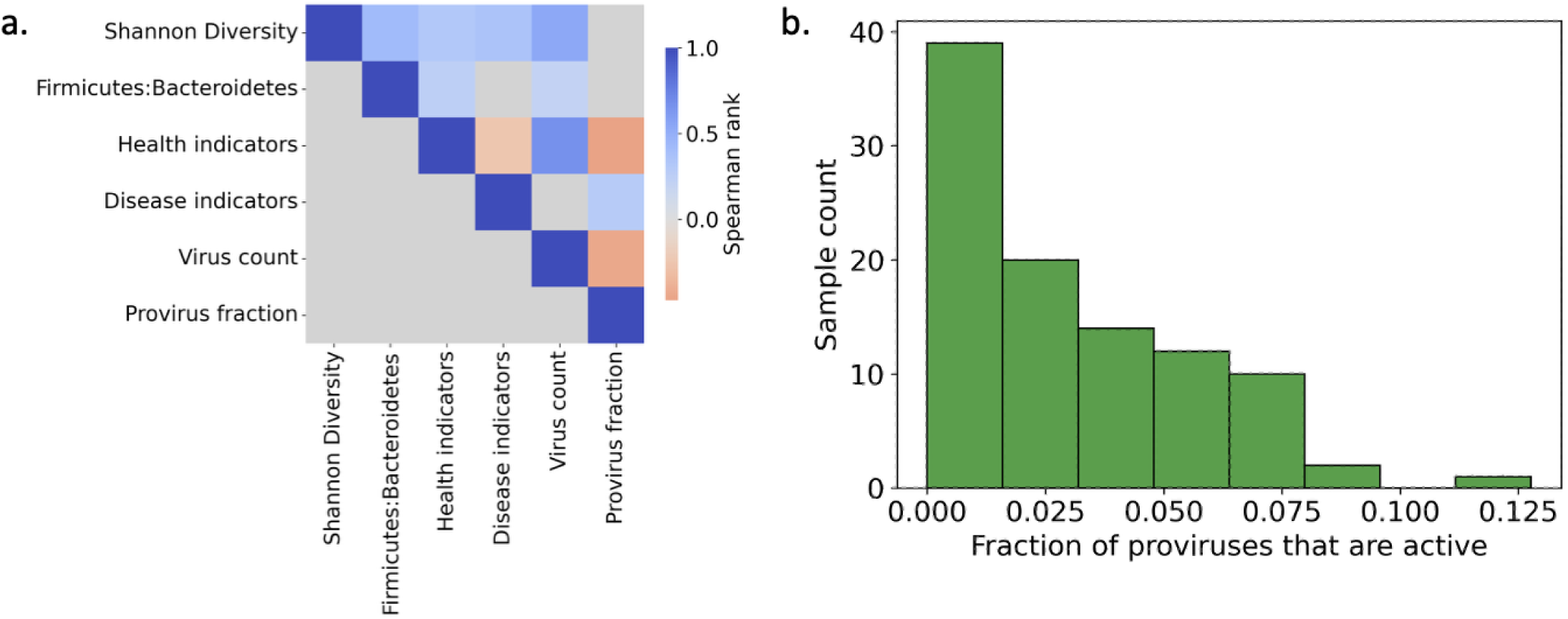
Further comparison of virome measures. (a) Comparison of viral population and bacterial population community-level metrics. (b) Histogram of active provirus fraction for SCD samples. Value of correlations is shown for comparisons with p *<* 0.05 otherwise comparison cell is gray.

**Supplemental Figure 2:**
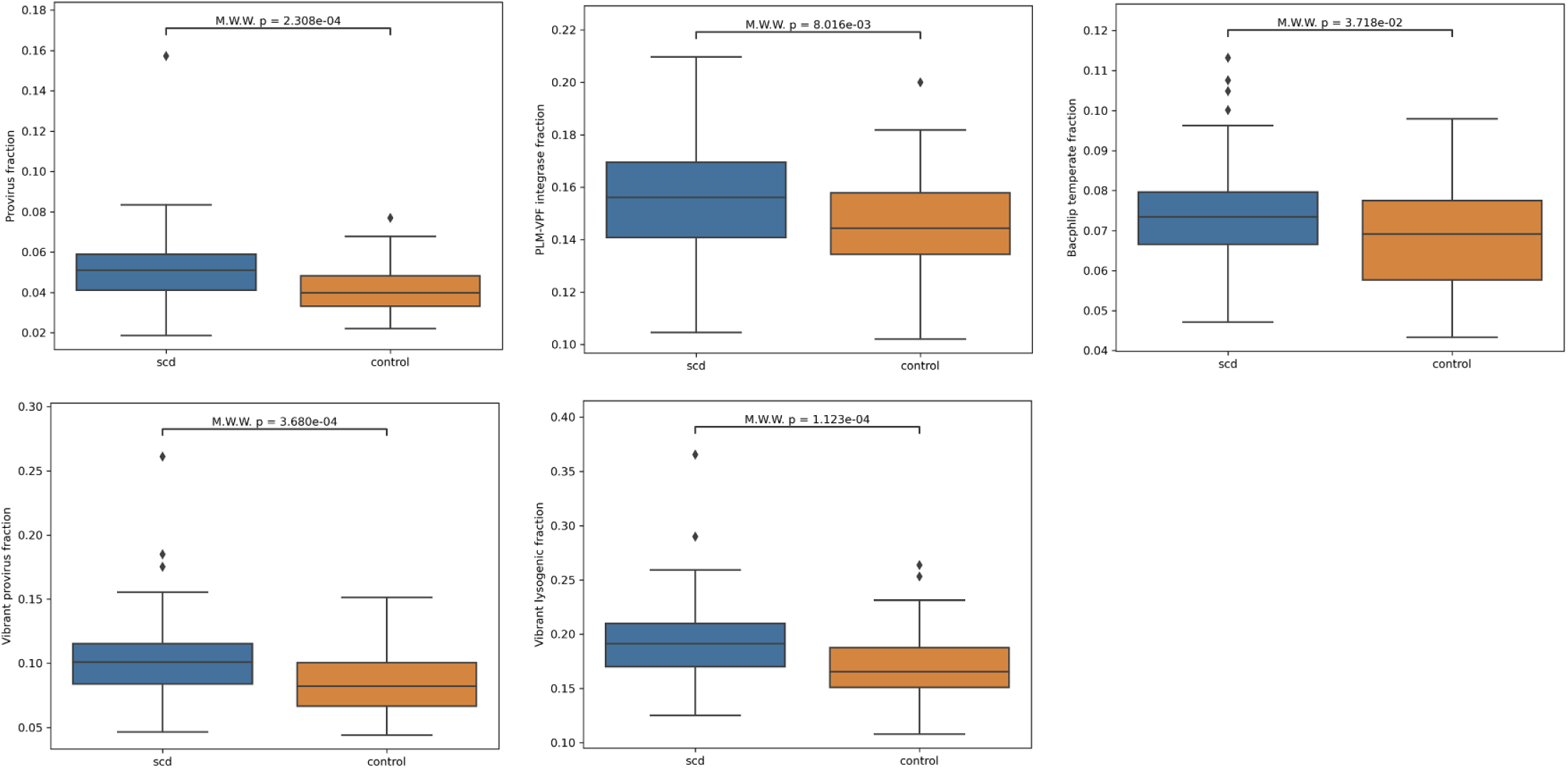
Enrichment in provirus or lysogenic virus prediction across multiple labeling methods. The determination of a lysogenic virus can be made by either looking for the virus sequence integrated into a host genome or by predicting that a viral sequence has lysogenic potential. We utilized multiple methods that rely on different approaches to determine that the provirus enrichment observed is robust to metho

**Supplemental Figure 3:**
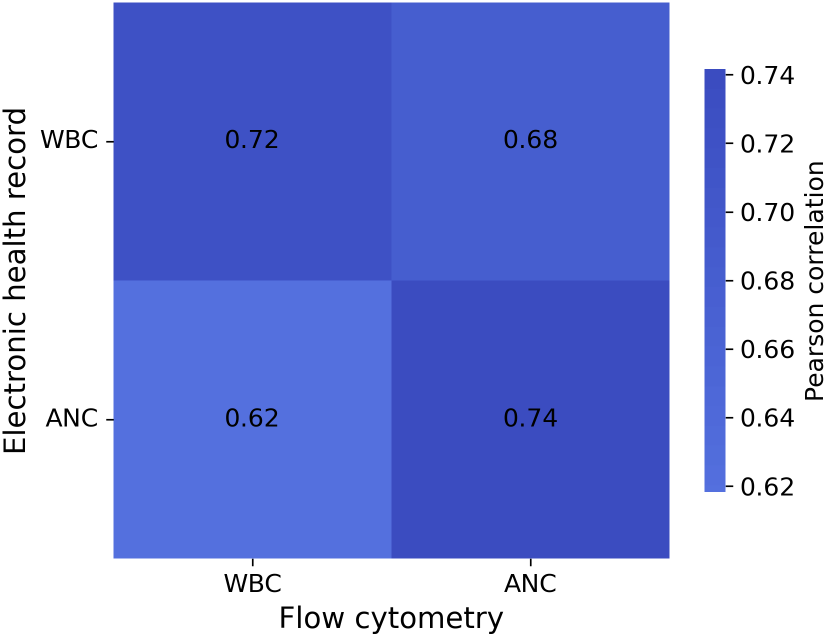
Correlation of flow cytometry and EHR measures of patient white blood cell populations. Correlation measured with Pearson correlation coefficient. Abbreviations: WBC-white blood cells, ANC-absolute neutrophil count.

**Supplemental Figure 4:**
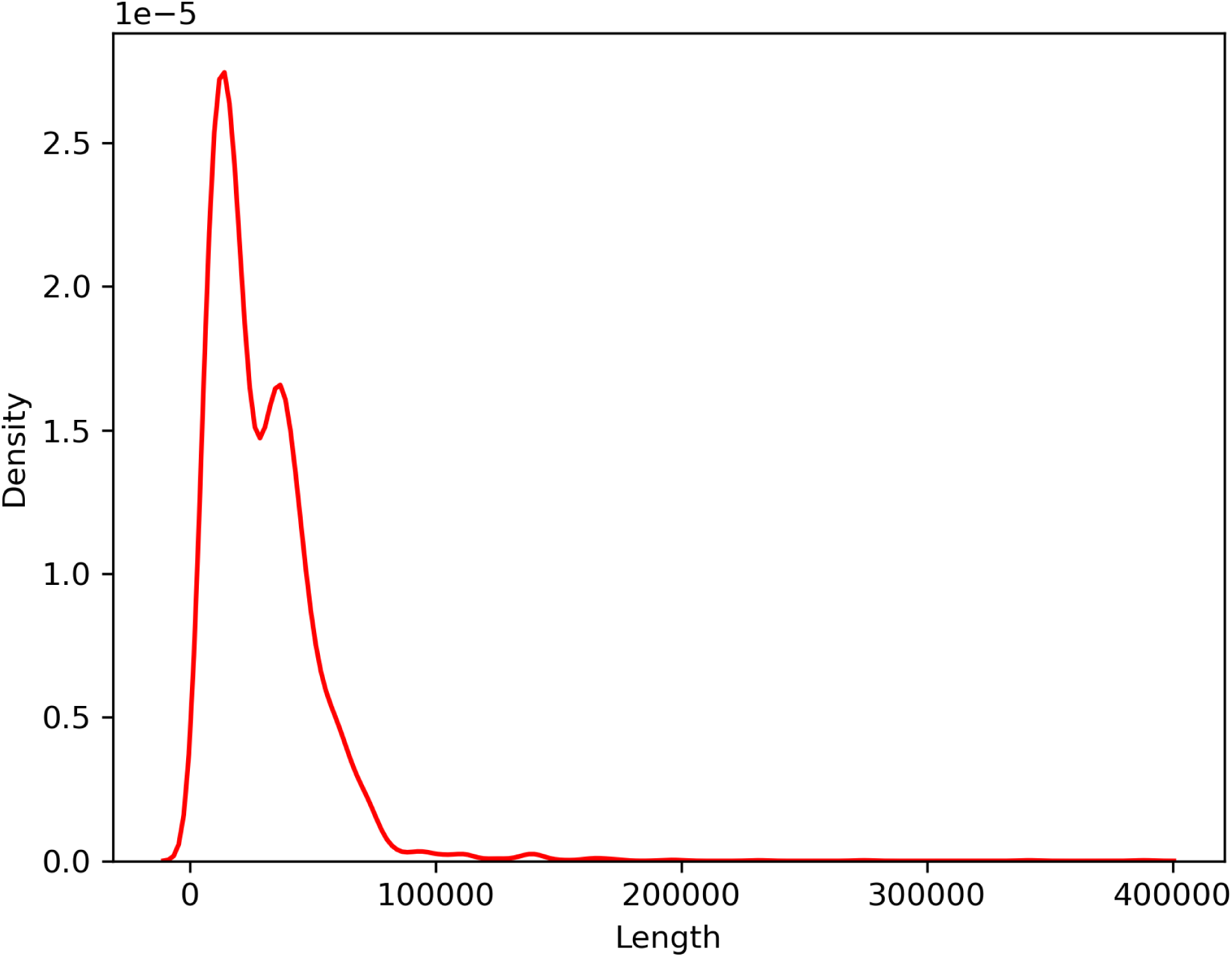
**Provirus length distribution with kernel density estimation.**

**Supplemental Figure 5:**
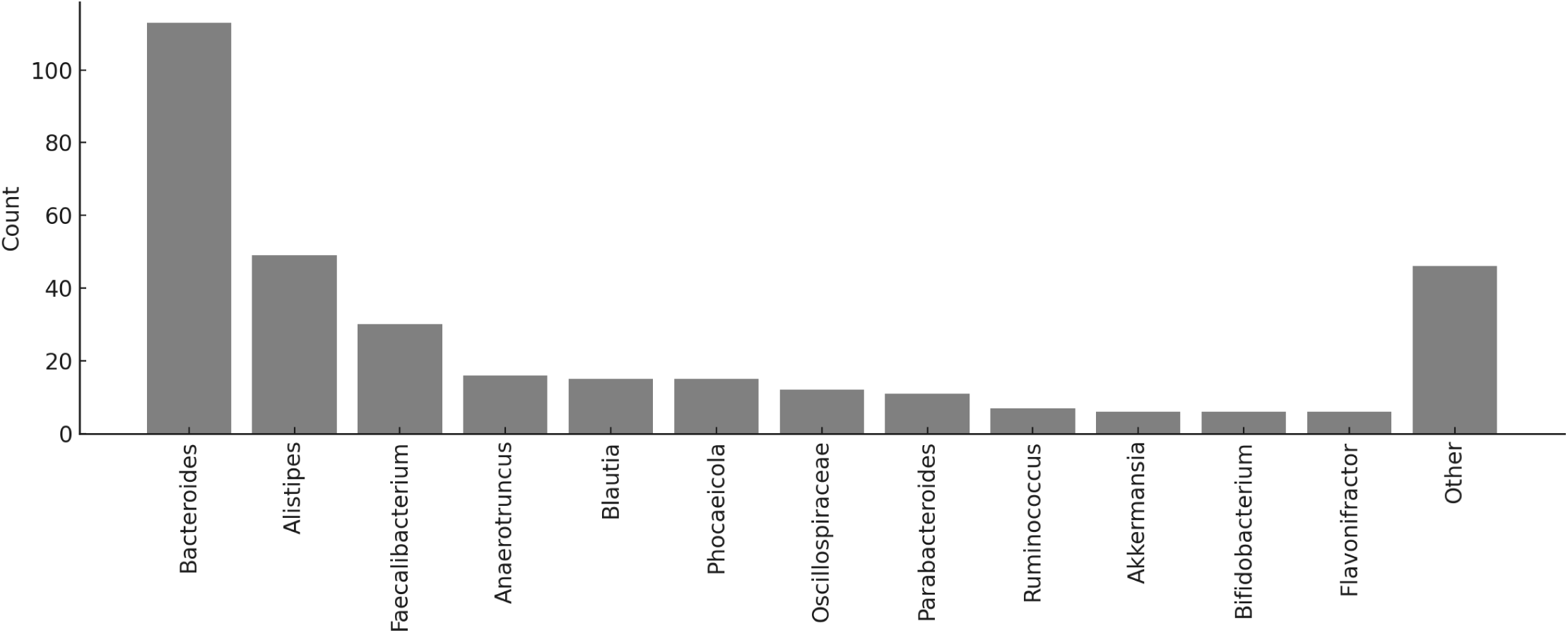
Prophage cluster sequence homology with bacterial hosts. Representative cluster sequences with blast sequence homology to bacterial species sequences were aggregated to the genus level. Count represents the number of clusters homologous to species in the genus.

**Supplemental Figure 6:**
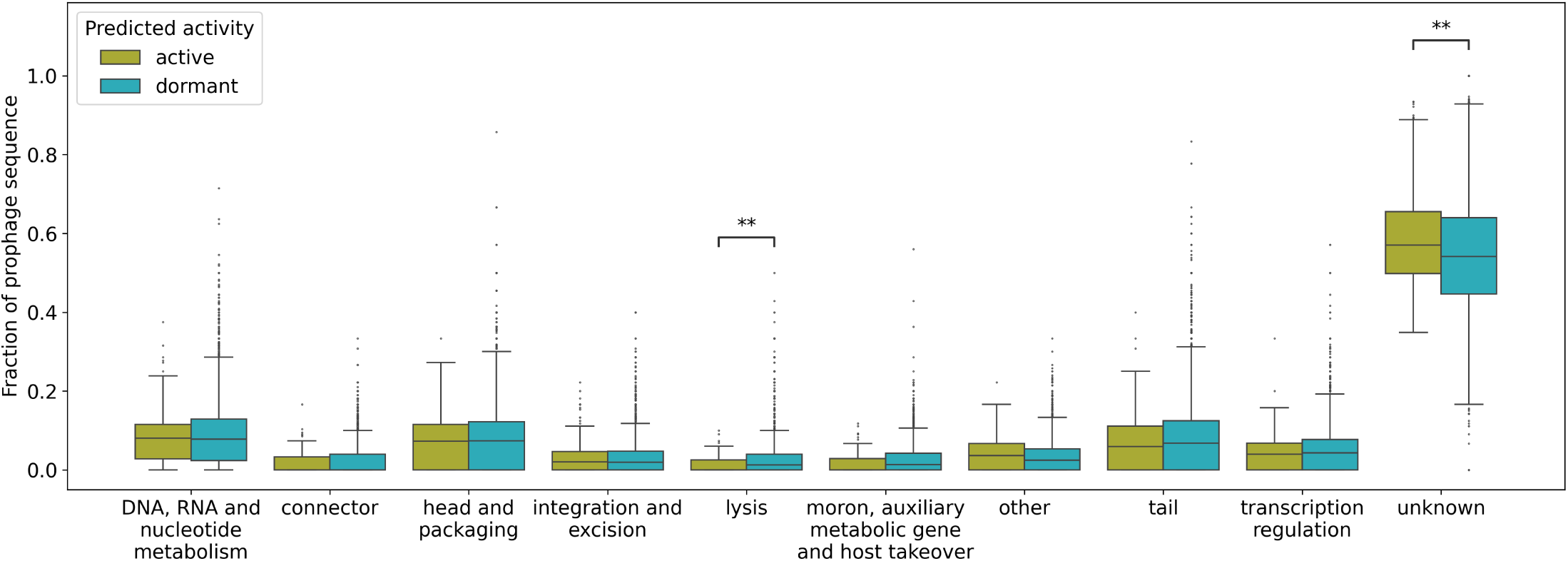
Genome content comparison between predicted active (n=136) and dormant (n=4739) prophages in the SCD gut microbiome. Significance tested with a Mann-Whitney-Wilcoxon test: * = 0.01 *<*= p *<* 0.05; ** = 0.001 *<* p *<*= 0.01.

**Supplemental Table 1:** Study participant demographics by cohort.

|  | SCD (n=98) | control (n=46) |
| --- | --- | --- |
| Sex (Female) [%] | 49 | 65 |
| Ethnicity (Non-Hispanic; Hispanic; Unknown) [%] | 78; 20; 2 | 74; 26; 0 |
| Race (Black; White; American Indian; Multiple race; Other/Unknown) [%] | 80; 1; 1; 1; 17 | 74; 0; 0; 0; 26 |
| Genotype [Hb type] | SS 92%,<br>Sβ-thalassemia 4%,<br>SCD with high HbF 4% | AS 61%,<br>AA 39% |
| Age [mean (standard deviation)] | 18.2 (12.8) | 18.6 (14.9) |

**Supplemental Table 2:** SCD patient (n=98) medical history.

| History (number with measure) | SCD patients [%] |
| --- | --- |
| Asthma (97) | 24 |
| Environmental allergies (97) | 43 |
| Stroke (96) | 7 |
| Acute chest syndrome (97) | 6 |
| Bacteremia (96) | 26 |
| History of bacteremia, meningitis, osteomyelitis, or urinary tract infection (96) | 30 |
| Acute care visits in past year (97) | 45 |
| ED visit in past year (97) | 35 |
| Pain admission in past year (97) | 22 |

**Supplemental Table 3:** SCD patient (n=98) treatment history.

| Treatment (number with measure) | SCD patients [%] |
| --- | --- |
| HU (98) | 62 |
| Folic acid (97) | 72 |
| Glutamine (97) | 16 |
| Transfusions in past year (97) | 37 |

**Supplemental Table 4:**
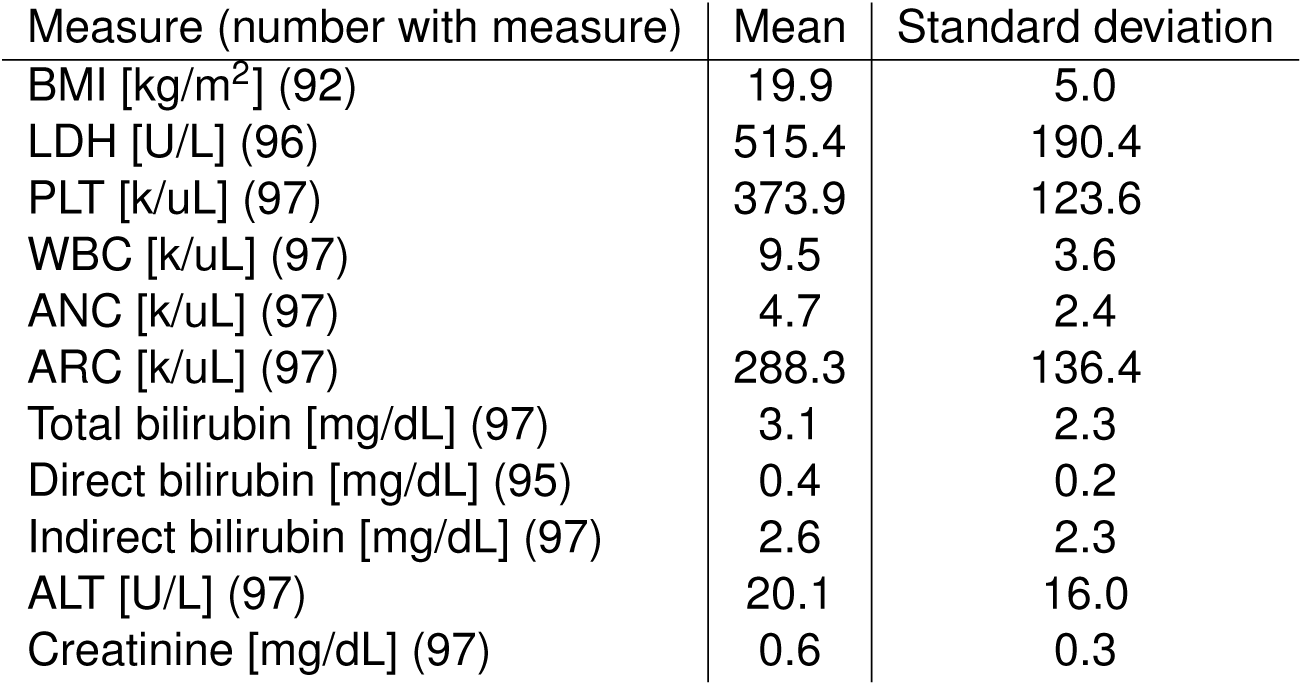
Clinical measures for SCD patients (n=98).

| Measure (number with measure) | Mean | Standard deviation |
| --- | --- | --- |
| BMI [kg/m<sup>2</sup>] (92) | 19.9 | 5.0 |
| LDH [U/L] (96) | 515.4 | 190.4 |
| PLT [k/uL] (97) | 373.9 | 123.6 |
| WBC [k/uL] (97) | 9.5 | 3.6 |
| ANC [k/uL] (97) | 4.7 | 2.4 |
| ARC [k/uL] (97) | 288.3 | 136.4 |
| Total bilirubin [mg/dL] (97) | 3.1 | 2.3 |
| Direct bilirubin [mg/dL] (95) | 0.4 | 0.2 |
| Indirect bilirubin [mg/dL] (97) | 2.6 | 2.3 |
| ALT [U/L] (97) | 20.1 | 16.0 |
| Creatinine [mg/dL] (97) | 0.6 | 0.3 |

**Supplemental Table 5:** Molecular inflammatory measures for SCD patients (n=98).

| Measure (number with measure) | Mean | Standard deviation | Fraction above lower limit of detection |
| --- | --- | --- | --- |
| *Cytokines* |  |  |  |
| G-CSF [pg/mL] (70) | 18.6 | 31.9 | 36% |
| IFNγ [pg/mL] (85) | 20.4 | 31.7 | 62% |
| IL-1α [pg/mL] (87) | 68.4 | 122.9 | 99% |
| IL-1β [pg/mL] (90) | 4.1 | 8.1 | 54% |
| IL-6 [pg/mL] (74) | 20.5 | 33.7 | 99% |
| IL-10 [pg/mL] (95) | 33.6 | 82.9 | 73% |
| IL-17A [pg/mL] (91) | 5.1 | 8.5 | 58% |
| TNF-α [pg/mL] (67) | 4.5 | 5.1 | 73% |
| *Chemokines* |  |  |  |
| IL-8 [pg/mL] (94) | 49.1 | 104.6 | 88% |
| IP-10 [pg/mL] (91) | 329.5 | 264.8 | 99% |
| MCP-1 [pg/mL] (92) | 67.5 | 52.4 | 99% |
| MIP-1 [pg/mL] (92) | 18.2 | 9.9 | 98% |

**Supplemental Table 6:** Analysis of genotype influence on significant microbiome and virome features.

| Feature | HbAA *n* | HbAS *n* | HbAA median | HbAS median | *p*-value | BH *q*-value |
| --- | --- | --- | --- | --- | --- | --- |
| F:B ratio | 18 | 28 | 2.228 | 2.215 | 0.328 | 0.437 |
| Shannon diversity | 18 | 28 | 3.155 | 3.117 | 0.536 | 0.536 |
| Provirus fraction | 18 | 28 | 0.0435 | 0.0375 | 0.166 | 0.333 |
| Virus count | 18 | 28 | 1157.5 | 1374.5 | 0.153 | 0.333 |

**Supplemental Table 7:** Testing associations of significant microbiome and virome features with treatment exposure or disease severity.

